# Increased Expression of Chondroitin Sulfotransferases following AngII may Contribute to Pathophysiology Underlying Covid-19 Respiratory Failure: Impact may be Exacerbated by Decline in Arylsulfatase B Activity

**DOI:** 10.1101/2020.06.25.171975

**Authors:** Sumit Bhattacharyya, Kumar Kotlo, Joanne K. Tobacman

## Abstract

The precise mechanisms by which Covid-19 infection leads to hypoxia and respiratory failure have not yet been elucidated. Interactions between sulfated glycosaminoglycans (GAGs) and the SARS-CoV-2 spike glycoprotein have been identified as participating in viral adherence and infectivity. The spike glycoprotein binds to respiratory epithelium through the angiotensin converting enzyme 2 (ACE2) receptor, which endogenously interacts with Angiotensin (Ang) II to yield Angiotensin 1-7. In this report, we show that stimulation of human vascular smooth muscle cells by Ang II leads to increased mRNA expression of two chondroitin sulfotransferases (CHST11 and CHST15), which are required for synthesis of chondroitin 4-sulfate (C4S) and chondroitin 4,6-disulfate (CSE), respectively. Also, increased total sulfated GAGs, increased sulfotransferase activity, and increased expression of the proteoglycans biglycan, syndecan, perlecan, and versican followed treatment by Ang II. Candesartan, an Angiotensin II receptor blocker (Arb), largely, but incompletely, inhibited these increases, and the differences from baseline remained significant. These results suggest that another effect of Ang II also contributes to the increased expression of chondroitin sulfotransferases, total sulfated GAGs, and proteoglycans. We hypothesize that activation of ACE2 may contribute to these increases and suggest that the SARS-CoV-2 spike glycoprotein interaction with ACE2 may also increase chondroitin sulfotransferases, sulfated GAGs, and proteoglycans and thereby contribute to viral adherence to bronchioalveolar cells and to respiratory compromise in SARS-CoV-2 infection.

## Introduction

The COVID-19 pandemic presents unique challenges, as the medical and scientific communities develop new therapies, new diagnostic tests, and new insights into the mechanisms by which the SARS-CoV-2 virus acts [1,2]. The angiotensin converting enzyme 2 (ACE2; Gene ID=59272) is the target for the spike glycoprotein receptor binding domain of the SARS-CoV-2 virus [3,4]. ACE2 is also the enzyme which converts Angiotensin (Ang) II to Ang (1-7), following the conversion of Ang I to the octapeptide Ang II by the loss of two amino acids. Physiologically, the vasodilator Ang (1-7) counteracts the vasoconstrictor effect of Ang II. The impact of the commonly used anti-hypertensive medications, angiotensin 1 converting enzyme inhibitors (ACEI) and angiotensin II receptor blockers (Arb), on Covid-19 disease have been a subject of intense discussion [5–7]. The experiments presented in this report were performed to detect effects of activation of the renin-angiotensin system (RAS) on chondroitin sulfates and associated proteoglycans, since these are largely unexplored. The experiments were performed to consider how Ang II exposure and inhibition by Arb treatment affected the expression of chondroitin sulfotransferases and associated proteoglycans in human vascular smooth muscle cells. In the setting of the Covid-19 pandemic, the experimental findings have been considered in the context of how interaction with ACE2 may contribute to the underlying pathophysiology of Covid-19 infection, in relation to potential effects on chondroitin sulfotransferases and chondroitin sulfates.

Data are presented which show increased expression of the chondroitin sulfotransferases CHST15 (carbohydrate sulfotransferase 15; N-acetylgalactosamine 4-sulfate 6-*O*-sulfotransferase; GalNAc4S-6ST) and CHST11 (carbohydrate sulfotransferase 11; chondroitin 4-*O*-sulfotransferase 1) in human vascular smooth muscle cells, following stimulation by exogenous Angiotensin (Ang) II. CHST15, which is also known as BRAG (B-cell Recombination Activating Gene), is required for the synthesis of chondroitin sulfate E (CSE; [GlcA-GalNAc-4S,6S]_n_, in which S corresponds to sulfate) from chondroitin 4-sulfate (C4S; [GlcA-GalNAc-4S]_n_). CSE is composed of alternating β-1,4- and β-1,3-linked D-glucuronate and D-*N*-acetylgalactosamine-4S,6S residues, and is found throughout human tissues. Increases in CHST15 or in CSE have been recognized in malignant cells and tissues and in models of tumor progression and tissue remodeling, including in pulmonary and cardiac tissues [8–10]. CHST11 is required for the synthesis of C4S, which is widely distributed throughout mammalian tissues. Increases in C4S are pathognomonic of congenital Mucopolysaccharidosis (MPS) VI, due to inherited mutations of the enzyme arylsulfatase B (ARSB; N-acetylgalactosamine-4-sulfatase) [11,12]. ARSB removes the 4-sulfate group at the non-reducing end of C4S and is required for degradation of C4S and of CSE. Accumulation of chondroitin sulfates contributes to pulmonary dysfunction in the mucopolysaccharidoses, cystic fibrosis, and lung diseases [11–14]. Reduced ARSB activity in association with increased chondroitin 4-sulfation has been implicated in aberrant cell signaling and transcriptional events in mammalian cells [15–18].

Earlier reports described the interaction of coronaviruses with the sulfated glycosaminoglycans (GAGs) heparin and heparan sulfate [19,20]. Recent studies have implicated these GAGs in viral attachment of the receptor binding domain (RBD) of the spike glycoprotein to the angiotensin converting enzyme 2 (ACE2) receptor in mammalian cells [21–23]. A galectin-like fold has been identified in the N-terminal domain of the spike protein of SARS-CoV-2, resembling similar structures in other coronaviruses [24–28]. The presence of this fold suggests potential binding with cell-based galactosides, such as those of chondroitin sulfates, which are the preferred binding partners of galectins.

The studies presented in this report demonstrate several findings that are pertinent to mechanisms of how SARS-CoV-2 infection might lead to respiratory insufficiency due to effects of chondroitin sulfates and their associated proteoglycans. These effects do not preclude binding interactions between heparin or heparan sulfate and the spike glycoprotein, but suggest potential for cell membrane attachment by other spike protein-chondroitin sulfate interactions. In addition, the effect of decline in activity of the enzyme arylsulfatase B (ARSB; N-acetylgalactosamine-4-sulfatase), in association with increases in CHST11 and CHST15 and increases in C4S and CSE, on Covid-19 disease is considered. Decline in ARSB has been associated previously with increased mRNA and protein expression of Interleukin-6, a known mediator of cytokine storm in Covid-19 infection [29,30]. The proposed mechanisms involving chondroitin sulfotransferases, chondroitin sulfates, and ARSB, may contribute to the underlying pathophysiology of Covid-19 infection, and attention to these pathways may help to design new approaches to reduce d morbidity and mortality.

## Materials and Methods

### Human cell samples

Human aortic smooth muscle cells (Lifeline Technology, Oceanside, CA, USA) were maintained in DMEM supplemented with 10% fetal bovine serum at 37°C in a humidified atmosphere with 5% CO_2_. Cells were passaged twice, then grown in multiwell culture plates to 75% confluency, and media was changed to DMEM without serum for 24 h to maintain quiescence. Cells were then treated with Angiotensin II (Ang II; 1 μM for 24 h; Sigma-Aldrich, St. Louis, MO), candesartan, an angiotensin II receptor blocker (Arb; 10 μM for 24h; Sigma-Aldrich), or their combination for 24 h, harvested and frozen for further experiments.

Circulating leukocytes were isolated from whole blood samples obtained from patients aged 2-17, followed at Rush University Medical Center (RUMC) for cystic fibrosis, asthma, or other conditions under a protocol approved by RUMC and the University of Illinois at Chicago (UIC) Institutional Review Boards [29]. Patients 8 years of age and older signed assent forms, and parents of all subjects signed consent forms, permitting collection of peripheral blood samples. Control subjects were children with a known, non-respiratory, underlying diagnosis or children with no known illness. Venous blood was collected in citrated tubes and processed within 2 h of collection. Subjects had no acute illness at the time of blood collection. Blood samples were identified with a study number, and a registry linked patient clinical data with study identifier. Polymorphonuclear (PMN) and mononuclear (MC; monocytes and lymphocytes) white blood cells were collected separately from the whole blood samples by the PolymorphprepTM kit (AXIS-SHIELD, Oslo, Norway) [29]. Cells were stored at −80°C until further experiments.

### Measurement of arylsulfatase B activity

ARSB activity was measured in the leukocyte samples using a fluorometric assay, following a standard protocol with 20 μl of homogenate in ddH_2_O and 80 μl of assay buffer (0.05M Na-acetate buffer with 20mmol barium acetate /l Na-acetate, pH 5.6) with 100 μl of substrate [5mM 4-methylumbelliferone sulfate (MUS) in assay buffer]. Materials were combined in microplate wells, and the microplate was incubated for 30 min at 37°C. The reaction was stopped by adding 150 μl of stop buffer (glycine–carbonate buffer, pH 10.7). Fluorescence was measured at 360 nm (excitation) and 465 nm (emission). ARSB activity was expressed as nmol/mg protein/h. and was derived from a standard curve prepared with known quantities of 4-methylumbelliferone at pH 5.6 [31,32].

### Measurement of Interleukin-6 by ELISA

Interleukin (IL)-6 in the patient plasma samples was measured by the Quantikine ELISA kit for human IL-6 (R&D Systems, Minneapolis, MN), as previously [29]. IL-6 in the samples was captured into the wells of a microtiter plate pre-coated with specific anti-IL-6 monoclonal antibody, and the immobilized IL-6 was detected by a biotinylated second antibody and streptavidin-horseradish peroxidase (HRP)-conjugate with the chromogenic substrate hydrogen peroxide/ tetramethylbenzidine (TMB). Color intensity was read at 450 nm with a reference filter of 570 nm in an ELISA plate reader (FLUOstar, BMG LABTECH, Inc., Cary, NC). IL-6 concentrations were extrapolated from a standard curve plotted using known concentrations of IL-6, expressed as picograms per milliliter plasma (pg/ml).

### QRT-PCR for sulfotransferases and proteoglycans

QRT-PCR was performed using established techniques and primers identified using Primer 3 software [33,34]. Primers were:

CHST11 human (NM_018413) Forward: 5’-GTTGGCAGAAGAAGCAGAGG-3’ and Reverse: 5’-GACATAGAGGAGGGCAAGGA-3’;
CHST15 human (NM_015892) Forward: 5’-ACTGAAGGGAACGAAAACTGG-3’ and Reverse: 5’-CCGTAATGGAAAGGTGATGAG-3’.
ARSB (NM_000046) Forward: 5’-AGACTTTGGCAGGGGGTAAT-3’ and Reverse: 5’-CAGCCAGTCAGAGATGTGGA-3’.
Biglycan (BC002416.2) Forward: 5’-ACTGGAGAACAGTGGCTTTGA-3’ and Reverse: 5’-ATCATCCTGATCTGGTTGTGG-3’.
CSPG4 (NM_001897) Forward: 5’-CTTCAACTACAGGGCACAAGG-3’ and Reverse: 5’-AGGACATTGGTGAGGACAGG-3’;
Perlecan (NM_001291860) Forward: 5’-CCTTGCTCAGAATGCACTAGG-3’ and Reverse: 5’-TGATGAGTGTGTCACCTTCCA-3’;
Syndecan-1 (NM_002997) Forward: 5’-CTCTGTGCCTTCGTCTTTCC-3’ and Reverse: 5’-CCACCTTCCTTTGCCATTT-3’;
TPST1 (NM_003596.3) Forward: 5’-GACCTCAAAGCCAACAAAACC-3’ and Reverse: 5’-TCAGTAACACCAGCCTCATCC-3’;
Versican (NC_000005.10) Forward: 5’-CCACTCTGTTTTCTCCCCATT-3’ and Reverse: 5’-ATCCCTTTGTGCCCTTTTTC-3’.

Relative expression was calculated by standard methods comparing treated and control cells using GAPDH as housekeeping gene [15].

### Measurement of total sulfated glycosaminoglycans and sulfotransferase activity

Measurement of total sulfated glycosaminoglycans (GAGs) was performed as previously described [17]. Briefly, the BlyscanTM assay kit (Biocolor Ltd, Newtownabbey, N. Ireland) was used for detection of the sulfated GAG, based on the reaction of 1,9-dimethylmethylene blue with the sulfated oligosaccharides in the GAG chains.

Sulfotransferase activity was determined using the Universal Sulfotransferase Activity kit (R&D, Minneapolis, MN), which assesses the sulfotransferase activity of all the sulfotransferases in the cells or tissue tested [17]. 3′-Phosphoadenosine-5′-phosphosulfate (PAPS) is the sulfate donor for the sulfotransferase reaction, and PAPS (10 μl, 1 mM), acceptor substrate (10 μl), and coupling phosphatase (3.5 μl, 100 ng/μl) were combined with 25 μl/well of the cell suspension in the wells of a microtiter plate. For negative controls, assay buffer was substituted for the cell suspension. The mixture was incubated for 20 min at 37°C, then 30 μl of malachite green was added and the mixture gently tapped. De-ionized water (100 μl) was added to each well, color was developed, and the optical density of each well was read and compared among the different wells. The amount of product formation was calculated using a phosphate standard curve, following subtraction of the negative control. The 3-inorganic phosphate released by the coupling phosphatase and detected by malachite green phosphate detection reagent, was proportional to the 3′-phosphoadenosine-5′-phosphate generated, thereby indicating the extent of the sulfotransferase reaction which utilized the sulfate group of PAPS.

### Statistical analysis

Data were analyzed using InStat software (GraphPad, San Diego, CA) or Excel spreadsheet software by unpaired t-tests, two-tailed, with adjustment for equal or unequal standard deviation, or by one-way ANOVA with Tukey-Kramer post-test. For control and AngII treatments, 9-15 biological samples were tested, with technical replicates of each determination. Plasma IL-6 determinations were performed with duplicate samples, using the average of two determinations. Spearman correlation coefficients was calculated using Instat. Box and whisker representation indicates mean value by “X”, median by a line in the box, 25% and 75% quartiles by box margins, minimum and maximum by lines extending from the box (whiskers), and outliers by points beyond the whiskers. P-values < 0.05 are considered statistically significant. Values are compared to untreated control and *** or ### represents p<0.001, ** or ## indicates p<0.01, and * or # indicates p<0.05. Study data of cultured aortic smooth muscle cells have not been reported previously. Patient data from peripheral blood leukocytes and serum were reported previously in a different format [29].

## Results

### Angiotensin II exposure increases mRNA expression of CHST11 and CHST15

Treatment of human aortic smooth muscle cells with Angiotensin (Ang) II led to marked increases in mRNA expression of CHST15 and CHST11. CHST15 increased to 3.1 ± 0.2 times the baseline level (p<0.001; n=9; unpaired t-test, two-tailed). Co-treatment with candesartan, an angiotensin (Ang)-II receptor blocker (Arb), did not completely inhibit the Ang II-induced increases in expression of CHST15. The difference between control and Ang II+Arb remained significant (p<0.001; n=6; unpaired t-test, two-tailed) (**Fig.1A,1B**). CHST11 mRNA expression increased to 2.4 ± 0.2 times the baseline level (p<0.001; n=15; unpaired t-test, two-tailed). The Ang II-induced increase was not completely inhibited by candesartan, and the difference between control and Ang II+Arb remained significant (p<0.001; n=6; unpaired t-test, two-tailed) (**Fig.1C,1D**).

**Fig. 1.**
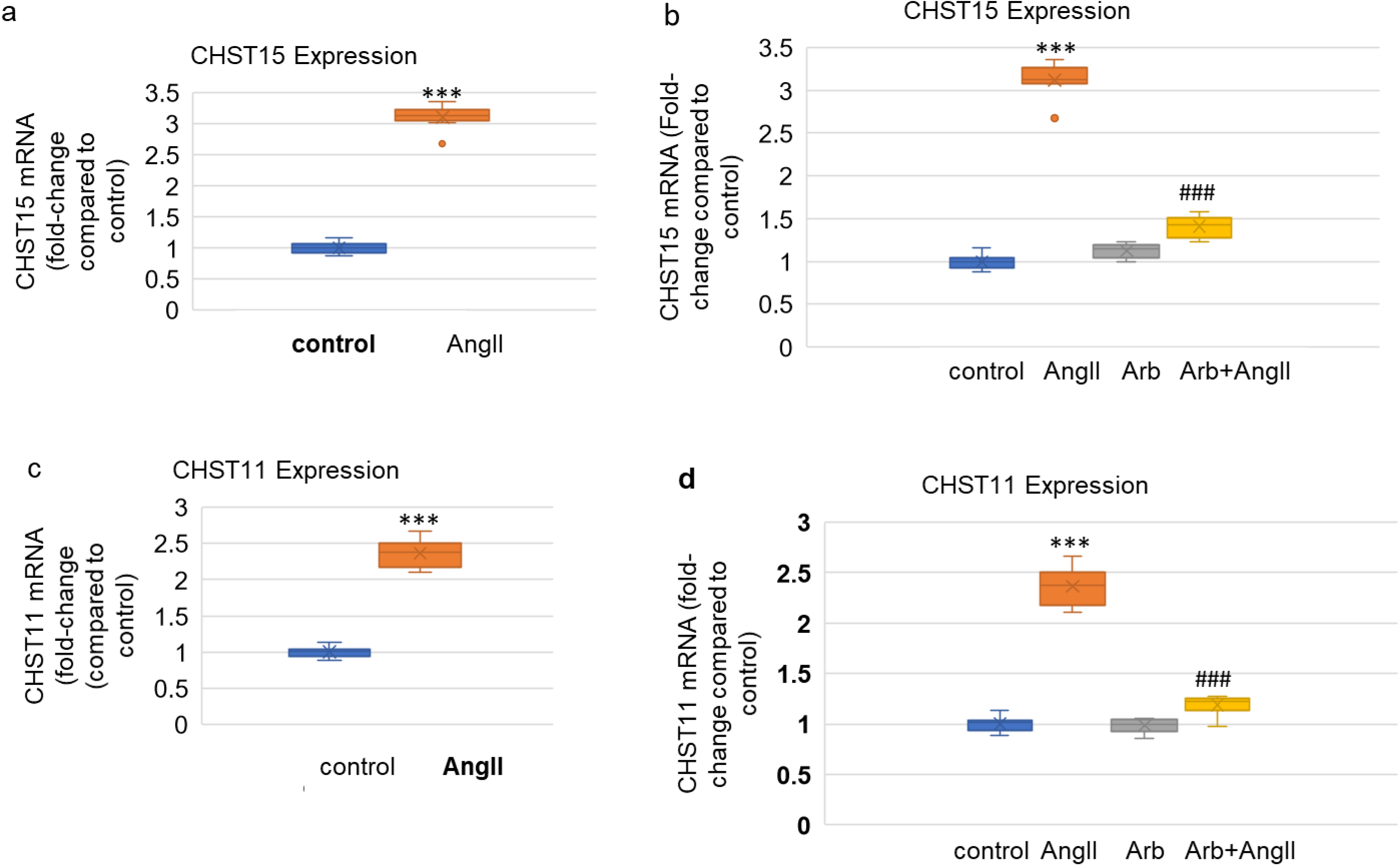
Increased Expression of CHST11 and CHST15 following AngII Exposure a. Following exposure to Ang II (1 μM x 24h), mRNA expression of CHST15 increased to 3.1 ± 0.2 times the baseline level (n=9, p<0.001, unpaired t-test, 2-tailed). Box and whisker representation indicates mean value by “X”, median by a line in the box, 25% and 75% quartiles by box margins, minimum and maximum by lines extending from the box (whiskers), and outliers by points beyond the whiskers. b. Treatment with the Ang II receptor blocker candesartan (10 μM x 24h) nearly completely blocked the Ang II-induced increase in CHST15 mRNA expression, but the difference between control and Ang II+Arb was still highly significant (n=6, p<0.001, unpaired t-test, two-tailed). c. Following exposure to Ang II (1 μM x 24h), mRNA expression of CHST11 increased to 2.4 ± 0.2 times the baseline level (n=15, p<0.001, unpaired t-test, two-tailed).d. Treatment with the Ang II receptor blocker candesartan (10 μM x 24h) nearly completely blocked the Ang II-induced increase in CHST11 mRNA expression, but the difference between control and Ang II+ARB was still highly significant (n=6, p<0.001, unpaired t-test, two-tailed). [Ang II=angiotensin II; ACE2=angiotensin converting enzyme 2; ARB=angiotensin II receptor blocker; CHST11=carbohydrate sulfotransferase 11=chondroitin 4-sulfotransferase 1; CHST15=carbohydrate sulfotransferase 15=N-acetylgalactosamine 4-sulfate 6-O-sulfotransferase]

Consistent with the effects of Ang II on CHST11 and CHST 15 expression, the total sulfated glycosaminoglycans (GAGs) increased (p<0.001; n=9; unpaired t-test, two-tailed). The increase was not completely inhibited by Arb treatment, and the difference between control and Ang II+Arb remained significant (p=0.048; n=3; unpaired t-test, two-tailed) (**Fig.2A,2B**). The measured sulfotransferase activity was also significantly increased following Ang II (p<0.001; n=9; unpaired t-test, two-tailed). The increase was largely inhibited by treatment with candesartan, but the difference between control and following treatment by Ang II+Arb remained significant (p=0.002, n=3; unpaired t-test, two-tailed) (**Fig.2C,2D**).

**Fig. 2.**
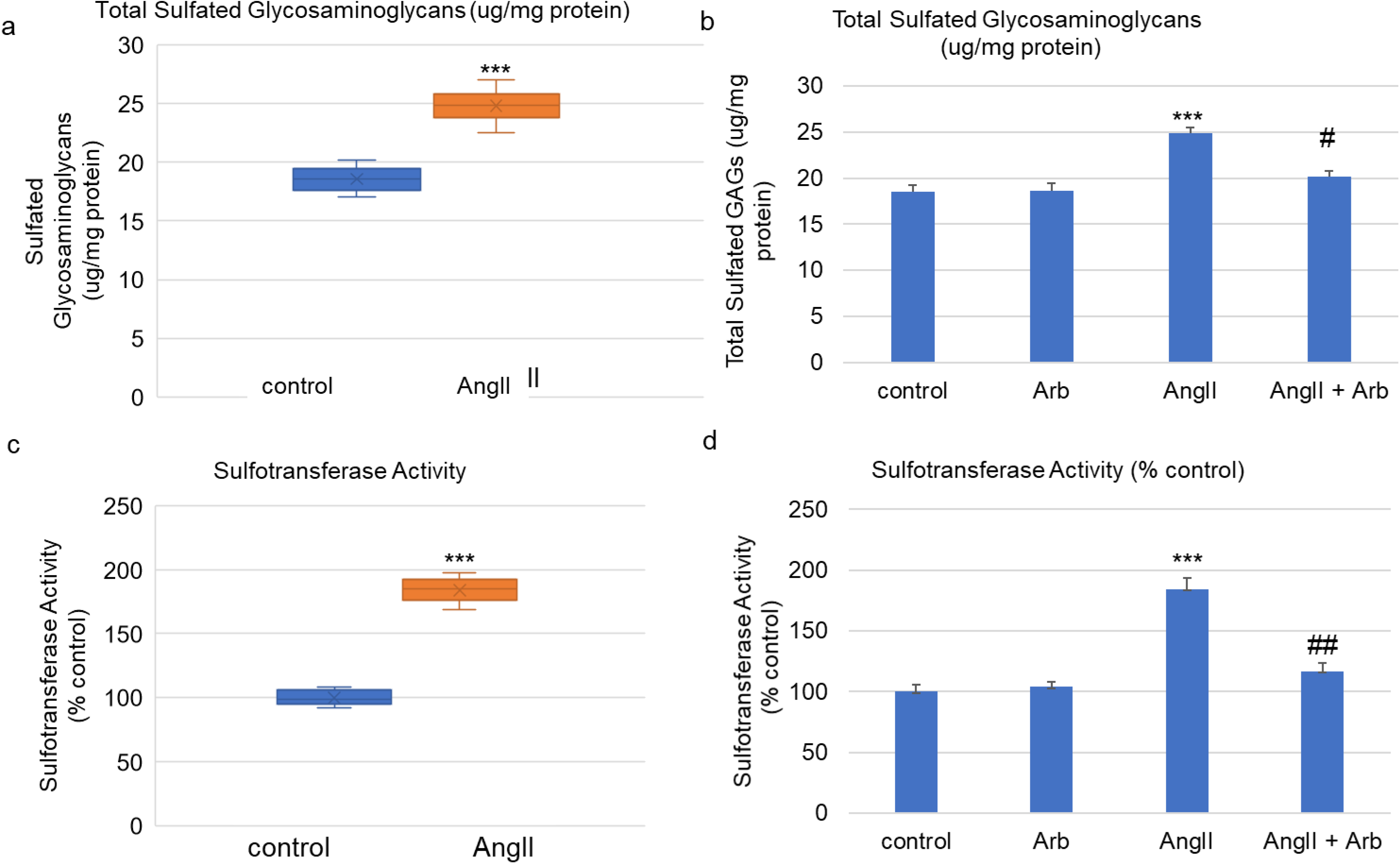
Sulfotransferase Activity and Total Sulfated Glycosaminoglycans a. Consistent with the increases in sulfotransferase activity and the increased expression of CHST11 and CHST15, the total sulfated glycosaminoglycans increased from 18.6 ± 1.1 μg/mg protein to 24.9 ± 1.5 μg/mg protein in the vascular smooth muscle cells following treatment by Ang II (p<0.001, n=9, unpaired t-test, two-tailed). Box and whisker representation indicates mean value by “X”, median by a line in the box, 25% and 75% quartiles by box margins, minimum and maximum by lines extending from the box (whiskers), and outliers by points beyond the whiskers. b. When treated with candesartan and Ang II, the peak total sulfated glycosaminoglycans declined by 4.8 μg/mg protein to 20.1 ± 0.7 μg/mg protein, reflecting an overall increase of 1.5 μg/ml. The difference between control value and post-treatment with Ang II and Arb remained statistically significant (p=0.048, n=3, unpaired t-test, two-tailed). c. Sulfotransferase activity increased to 184 ± 9% over the control level following Ang II (p<0.001, n=9, unpaired t-test, two-tailed). d. Sulfotransferase activity declined to 116 ± 7% of the control level when cells were treated with an Arb and Ang II. The difference between control value and post-treatment with Ang II and Arb remained significant (p=0.002, n=3, unpaired t-test, two-tailed). [Arb=angiotensin II receptor blocker; CHST11=carbohydrate sulfotransferase 11=chondroitin 4-sulfotransferase 1; CHST15=carbohydrate sulfotransferase 15=N-acetylgalactosamine 4-sulfate 6-O-sulfotransferase]

### Increased expression of chondroitin sulfate-associated proteoglycans

Following treatment with Ang II, mRNA expression of several proteoglycans which may have chondroitin sulfate attachments increased. These include increases from untreated control values for versican, syndecan-1, biglycan, and perlecan (p<0.001, n=6) (**Fig.3A**). In contrast to these increases, there was no increased expression of chondroitin sulfate proteoglycan (CSPG)4, tyrosylprotein sulfotransferase (TPST)1, or arylsulfatase B (ARSB) following Ang II exposure (data not shown). Treatment with candesartan largely inhibited the increases in expression of these proteoglycans, but the differences between control and Ang II + Arb remained significant (**Fig.3B**).

**Fig. 3.**
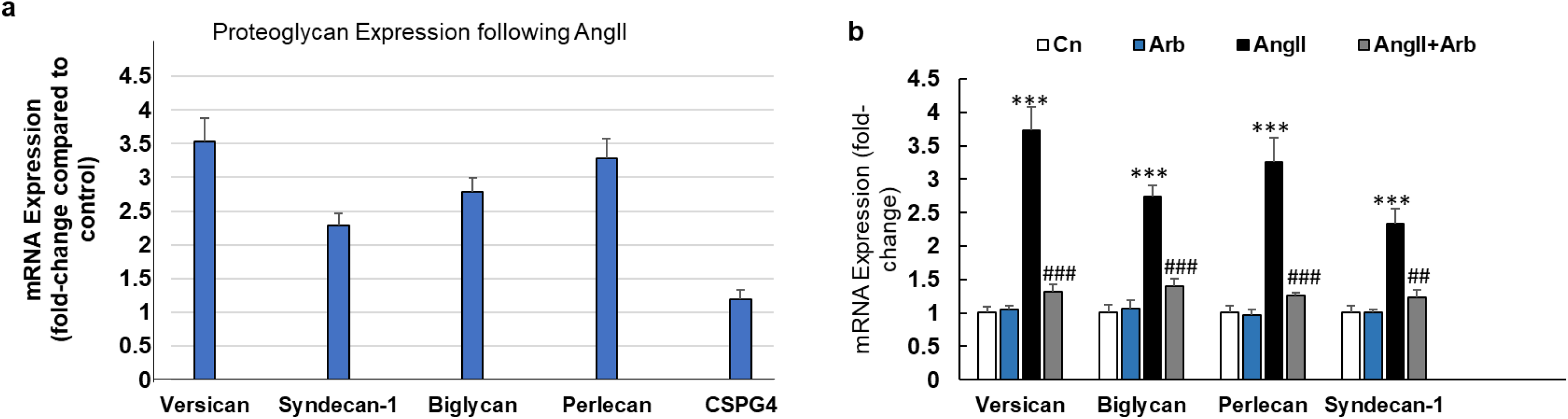
Impact of AngII and Blocker on Proteoglycan mRNA Expression a. mRNA expression of several proteoglycans is increased following treatment by Ang II (n=6). Increases were shown in versican, syndecan-1, biglycan, and perlecan, but not in CSPG4 or in TPST1 (not shown). b. Treatment with candesartan reduced, but did not eliminate, increases in the proteoglycans (n=6). [Cn=control; Arb=angiotensin II receptor blocker; Ang II=angiotensin II; CSPG4=chondroitin sulfate proteoglycan 4]

### Increase in IL-6 in association with decline in ARSB

In plasma from patients with cystic fibrosis or asthma, Interleukin-6 values were significantly greater in the plasma from the patients with cystic fibrosis and in patients with asthma, in contrast to the controls (p<0.001) (**Fig.4A**) [29]. Also, neutrophil Arylsulfatase B activity in CF and asthma was less compared to levels in neutrophils from healthy normal controls (p<0.001) (**Fig.4B**) [29]. An overall inverse relationship between ARSB and IL-6 in the control subjects and subjects with asthma or cystic fibrosis is apparent (Spearman r = −0.69) (**Fig.4C**).

**Fig. 4.**
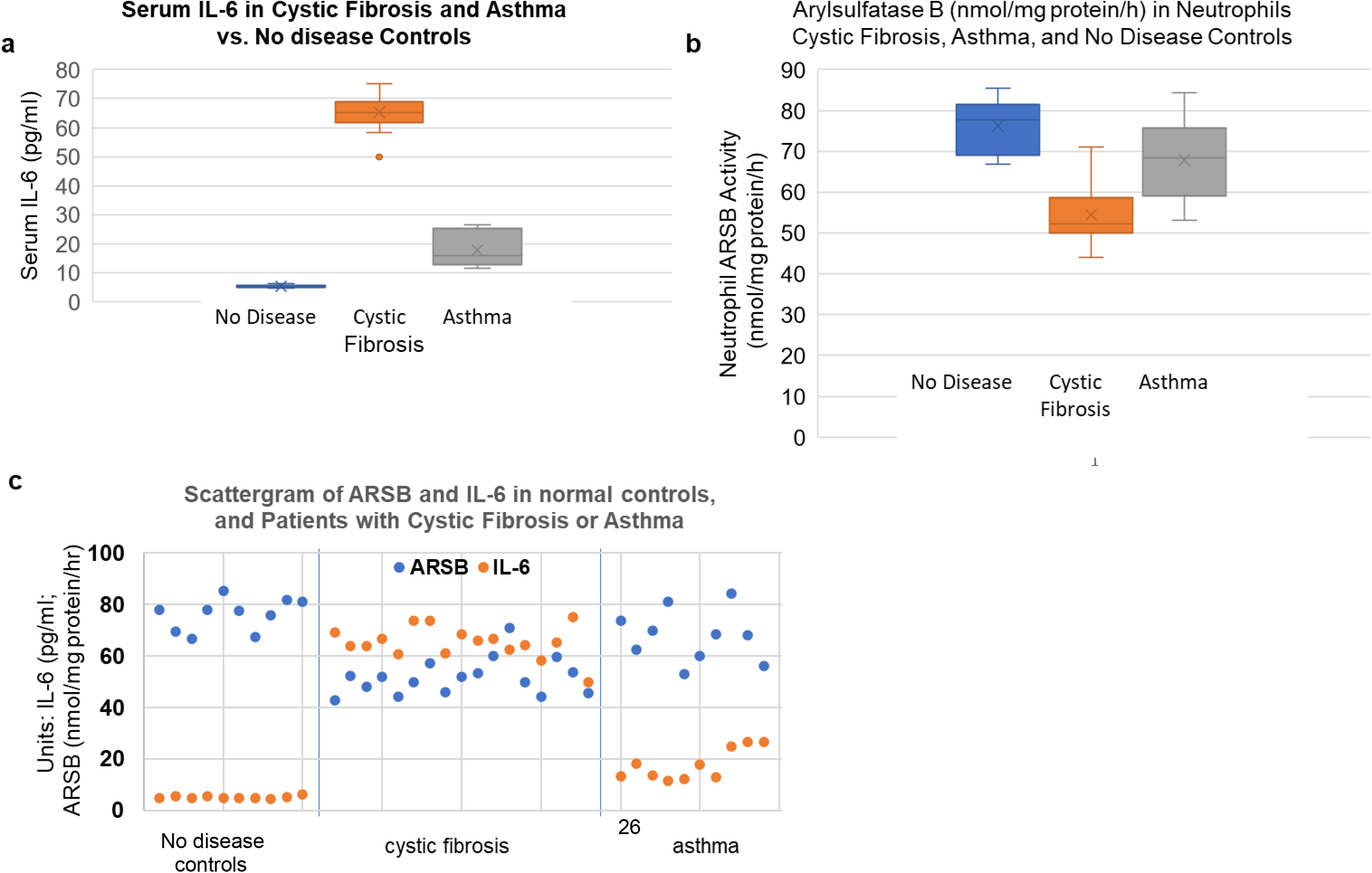
Inverse relationship between Interleukin-6 and Arylsulfatase B. a. Plasma Interleukin (IL)-6 levels were markedly increased in cystic fibrosis (n=17) compared to the normal controls (65.3 ± 6.2 pg/ml vs. 5.3 ± 0.5 pg/ml; n=10; p<0.001). Levels in patients with asthma (17.8 ± 6.1 pg/ml; n=10) were also significantly increased [29]. b. ARSB activity in the circulating neutrophils was markedly reduced in the cystic fibrosis patients (n=17), compared to the normal controls (n=10) (51.9 ± 7.2 vs. 76.2 ± 6.3 nmol/mg protein/h) and the patients with asthma (n=10) (67.8 ± 10.2 nmol/mg protein/h) [29]. c. There is an inverse relationship between plasma IL-6 levels and neutrophil ARSB activity (Spearman correlation coefficient r = −0.69). [ARSB=arylsulfatase B=N-acetylgalactosamine-4-sulfatase; IL-6 = interleukin-6]

### Overall schematic showing the potential impact of increase in sulfotransferases and of decline in chondroitin sulfatases in the pathophysiology of Covid-19

An overall representation of the proposed interactions between spike protein receptor binding with the ACE2 receptor indicates increased expression of CHST11 and CHST15 (**Fig. 5**). Increases in these sulfotransferases leads to increased C4S and CSE production, which accumulate and lead to hypoxia (↓pO_2_) and decline in ARSB activity. Decline in ARSB leads to increased expression of IL-6, thereby contributing to cytokine storm. When ARSB is reduced, the degradation of chondroitin sulfates is inhibited, further exacerbating effects of increased chondroitin sulfotransferase expression and chondroitin sulfate production initiated by ACE2 stimulation.

**Fig. 5.**
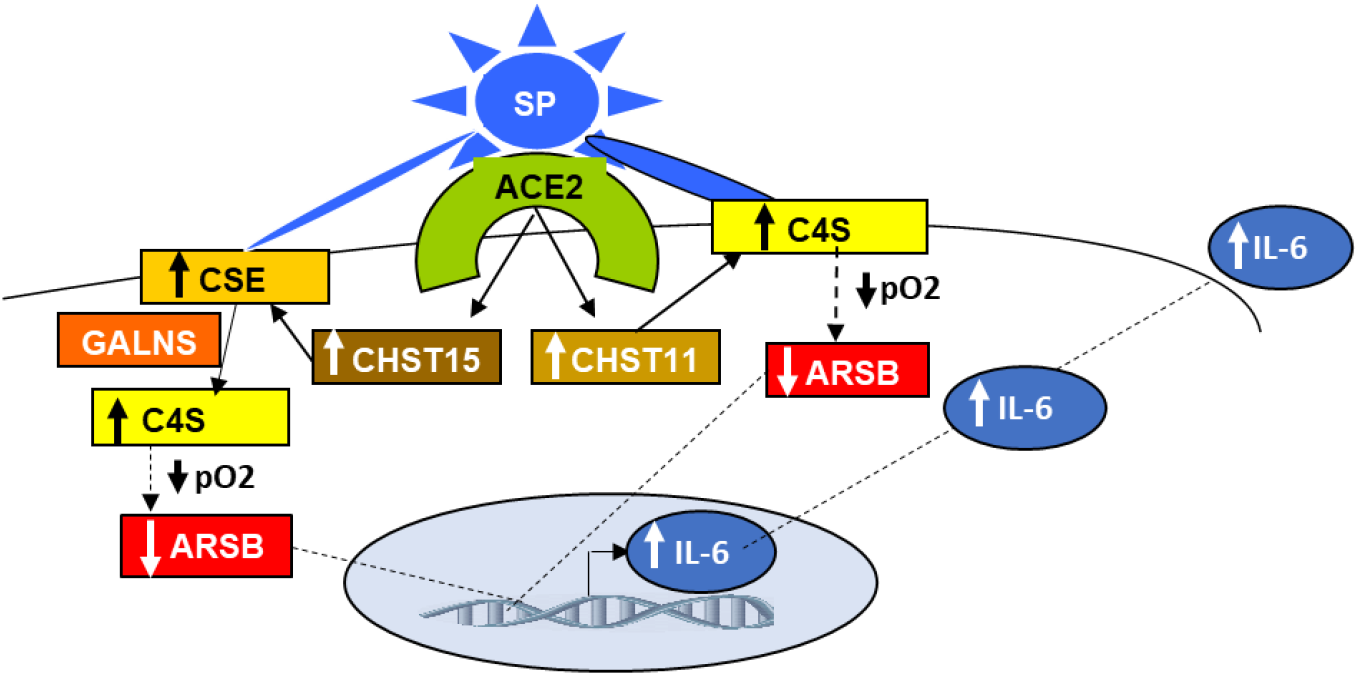
Schematic of Proposed Overall Pathway The overall pathway proposes that transcriptional events arise following stimulation of the ACE2 receptor lead to increased expression of CHST11 and CHST15. Increased sulfotransferase expression lead to increased production of C4S and CSE. CSE is converted to C4S by GALNS (N-acetylgalactosamine-6-sulfatase), producing C4S. The accumulation of sulfated glycosaminoglycans in the airways impairs oxygenation (↓O_2_), inhibiting the post-translational modification and activity of ARSB, leading to further accumulation of C4S, thereby aggravating respiratory insufficiency. Decline in ARSB also leads to increased expression of IL-6, which contributes to cytokine storm. [ACE2=angiotensin converting enzyme 2; ARSB=arylsulfatase B=N-acetylgalactosamine-4-sulfatase; CHST11=carbohydrate sulfotransferase 11=chondroitin 4-sulfotransferase 1; CHST15=carbohydrate sulfotransferase 15=N-acetylgalactosamine 4-sulfate 6-O-sulfotransferase=GalNAc4S-6ST; C4S=chondroitin 4-sulfate=[GlcA-GalNAc-4S]_n_ in which S is for sulfate; CSE=[GlcA-GalNAc-4S,6S]_n_, in which S is for sulfate; GALNS=N-acetylgalactosamine-6-sulfatase]

## Discussion

In the human aortic smooth muscle cells, exposure to Angiotensin (Ang) II markedly increased the mRNA expression of the chondroitin sulfotransferases CHST11 and CHST15. Sulfotransferase activity and total sulfated glycosaminoglycans, as well as the proteoglycans versican, perlecan, biglycan, and syndecan were also increased. Expression of TPST1, a tyrosylprotein sulfotransferase, chondroitin sulfate proteoglycan (CSPG)4, and arylsulfatase B was not increased. The angiotensin II receptor blocker (Arb) candesartan incompletely inhibited these increases, and increases remained significant. Since Ang II endogenously interacts with both the Ang II receptor (AT_1_) and the angiotensin-converting enzyme 2 (ACE2), the findings suggest either incomplete blockade of Ang II by candesartan or an additional effect of Ang II, perhaps mediated by interaction with ACE2. We propose consideration of effects mediated by ACE2, since these may be relevant to underlying mechanisms of SARS-CoV-2 pathophysiology.

Interactions between heparin and heparan sulfate and the receptor binding domain (RBD) of the spike glycoprotein do not preclude additional effects of chondroitin sulfates and proteoglycans in the pathophysiology of Covid-19 [19–23]. Identification of a galectin-like fold in the spike glycoprotein suggests that additional interactions with β-galactosides may contribute to binding of SARS-CoV-2 to cell surfaces [24–28]. Clausen *et al* reported that in A375 cells in which ACE2 was overexpressed, knockout of B4GALT7 (beta-1,4-galactosyltransferase 7), a gene required for proteoglycan and GAG assembly, reduced the binding of the spike glycoprotein [21]. This finding reflects a crucial role of glycosaminoglycans and proteoglycans in adherence of SARS-CoV-2 to human cells.

As in the mucopolysaccharidoses and in cystic fibrosis, accumulation of chondroitin sulfates in the lung is anticipated to impair airflow and oxygenation [11-13,32]. Prior work has shown a requirement of oxygen for post-translational modification and activation of arylsulfatase B (ARSB; N-acetylgalactosamine-4-sulfatase) [30,35]. Refractoriness to benefit of supplemental oxygen was demonstrated in patients with moderate chronic obstructive pulmonary disease who had mutations limiting ARSB expression [36]. In other studies, decline in ARSB increased IL-6 expression in bronchial epithelial cells [30], and plasma IL-6 was increased in patients with cystic fibrosis and asthma (**Fig.4**) [29]. The observed increases in CHST11 and CHST15 are anticipated to lead to increased C4S and CSE production, and these chondroitin sulfates are anticipated to accumulate further when ARSB activity is reduced. Increased chondroitin sulfate production by bronchial epithelial cells, leading to airway obstruction and reduced oxygenation, which lead to reduced ARSB activity and impaired degradation of chondroitin sulfates, may contribute to an ongoing cycle in Covid-19 culminating in refractory respiratory insufficiency (**Fig.5**).

In the future, it will be informative to develop these findings in bronchial epithelial cells and to quantify changes in chondroitin 4-sulfate, chondroitin sulfate E and total chondroitin sulfates. Ideally, these measurements should follow exposure of the BEC to spike glycoprotein or a modified coronavirus. At this time, we lack specific measurements of C4S, CSE, and ARSB activity in bronchial epithelial cells following Ang II or activation of ACE2 by the spike glycoprotein. Although lacking this information, the current findings suggest into how Covid-10 infection might evolve in a self-perpetuating cycle due to effects on chondroitin sulfotransferases and chondroitin sulfates and ARSB. The manifestations of lower ARSB may include: a) impact on viral binding to respiratory tract cells by the galectin-like fold of the spike glycoprotein when chondroitin sulfates are increased; b) impaired responsiveness to oxygen treatment; and c) enhanced IL-6 production contributing to cytokine storm. Recombinant human (rh) ARSB is used for replacement in Mucopolysaccharidosis VI (MPS VI), the inherited genetic deficiency of ARSB [11], and rhARSB may be a useful, new approach to refractory hypoxia in Covid-19 patients.

## Acknowledgment

The authors acknowledge the contributions of Robert Danziger, MD, MBA to vascular studies and the facilities of the Jesse Brown VAMC.

## Declarations

### Ethics approval

IRBs of the University of Illinois at Chicago and Rush University Medical Center approved the research which was previously reported.

### Consent to participate

Participants or their parents provided consent for participation.

### Consent for publication

Consent for publication was provided.

### Availability of data and material (data transparency)

All pertinent data are included in this publication and additional material can be obtained by communication with the authors.

### Code availability (software application or custom code)

NA

### Authors’ contributions

SB, KK, and JKT planned and performed the studies in the vascular smooth muscle cells. SB and JKT planned and performed the studies in the bronchial epithelial cells and the human blood samples. SB and JKT analyzed the data and JKT wrote the manuscript.

